# The complete genome sequence of the *Staphylococcus* bacteriophage Metroid

**DOI:** 10.1101/2020.05.01.072256

**Authors:** Adele Crane, Joy Abaidoo, Gabriella Beltran, Danielle Fry, Colleen Furey, Noe Green, Ravneet Johal, Bruno La Rosa, Catalina Lopez Jimenez, Linh Luong, Garett Maag, Jade Porche, Lauren Reyes, Aspen Robinson, Samantha Sabbara, Lucia Soto Herrera, Angelica Urquidez Negrete, Pauline Wilson, Kerry Geiler-Samerotte, Susanne P. Pfeifer

**Affiliations:** School of Life Sciences, Arizona State University, Tempe, USA; Center for Evolution and Medicine, Arizona State University, Tempe, USA; Center for Mechanisms of Evolution, Arizona State University, Tempe, USA

**Keywords:** bacteriophage, *Staphylococcus*, *Myoviridae*, genome assembly

## Abstract

Phages infecting bacteria of the genus *Staphylococcus* play an important role in their host’s ecology and evolution. On one hand, horizontal gene transfer from phage can encourage the rapid adaptation of pathogenic *Staphylococcus* enabling them to escape host immunity or access novel environments. On the other hand, lytic phages are promising agents for the treatment of bacterial infections, especially those resistant to antibiotics. As part of an ongoing effort to gain novel insights into bacteriophage diversity, we characterized the complete genome of the *Staphylococcus* bacteriophage Metroid, a cluster C phage with a genome size of 151kb, encompassing 254 predicted protein-coding genes as well as 4 tRNAs. A comparative genomic analysis highlights strong similarities – including a conservation of the lysis cassette – with other *Staphylococcus* cluster C1 bacteriophages, several of which were previously characterized for therapeutic applications.

## Introduction

Pathogens of the genus *Staphylococcus*, known for their ability to evade the human immune system, are an important public health concern causing a multitude of community-acquired infections ranging from food poisoning to skin lesions and life-threatening sepsis (Pollitt et al. 2018). As *Staphylococcus* largely reproduces clonally, much of the genetic diversity among strains stems from horizontal gene transfer through bacteriophages. Thereby, the acquisition of novel genes may not only aid adaptation of a bacterial strain to novel environments (Xia and Wolz 2014), but it can also increase pathogenicity. Bacteriophages play an important role in bacterial pathogenesis (Deghorain et al. 2012) as they encode for many known staphylococcal virulence factors (including the immune-modulator staphylokinase, as well as exfoliative Panton-Valentine leukocidin toxins; see review by Malachowa and DeLeo 2010). Moreover, bacteriophages can mediate the mobilization and transfer of genomic pathogenicity islands (Xia and Wolz 2014). On the other hand, virulent bacteriophages, which lyse their host cell after successful reproduction, also represent promising new avenues for the treatment of antibiotic-resistant *Staphylococcus* infections through phage therapy (Moller et al. 2019).

Approximately 10^30^ bacteriophages are estimated to exist on our planet (Rohwer 2003), however much of their diversity remains under-sampled and therefore uncharacterized. Several *Staphylococcus* phages (order: *Caudovirales*; *i.e.*, tailed dsDNA phages) have been isolated and sequenced (*e.g.,* Kwan et al. 2005; Deghorain et al. 2012; Olivera et al. 2019). Historically, *Staphylococcus* phages were grouped according to their lytic activity and serology; specifically, their reaction to (amongst others) polyclonal antiserum (Rountree 1949; Rippon 1952, 1956). In contrast, modern phage classification systems are based on either: 1) morphology (determined using transmission electron microscopy), categorizing *Myoviridae* (long, contractile tail; group A), *Siphoviridae* (long, non-contractile tail; group B), and *Podoviridae* (short tail; group C) (Ackermann 1975; Brandis and Lenz 1984); or 2) genome size, categorizing class I (<20kb), class II (~40kp), and class III (>125kb) (Kwan et al. 2005), with phages of like category generally being more closely related to one another (Kwan et al. 2005). More recently, in one of the largest *Staphylococcus* phage genomic studies published to date, Olivera et al. (2019) developed a comparative evolutionary approach to group *Staphylococcus* phages according to their gene content: cluster A (morphologically *Podoviridae;* genome size: 16-18kb), cluster B (a diverse cluster consisting of mostly temperate phages; genome size: 39-48kb), cluster C (morphologically *Myoviridae;* genome size: 127-152kb), and cluster D (morphologically *Siphoviridae;* genome size: 89-93kb). Based on predicted sequence similarities of protein families (phams), the authors further subdivided *Staphylococcus* phages into 27 subclusters (A1-A2, B1-B17, C1-C6, and D1-D2), members of which exhibit similar morphology and genomic features (*i.e.*, genome size, GC-content, and number of genes; Olivera et al. 2019). In contrast to the usually temperate *Siphoviridae*, most *Myoviridae* and *Podoviridae* experimentally characterized to date exhibit a lytic life cycle. Lytic phages destroy their host cells, making them interesting candidates for phage therapy (Xia and Wolz 2014).

Here, we report the complete genome sequence of the *Staphylococcus* bacteriophage Metroid, a *Myoviridae* sequenced as part of HHMI’s SEA-PHAGES program – an ongoing effort to systematically characterize bacteriophages and their relationship to their (often pathogenic) bacterial hosts. A comparative genomic analysis highlights strong similarities with other *Staphylococcus* cluster C1 bacteriophages, several of which were previously characterized for therapeutic applications (Vandersteegen et al. 2011; Gill 2014; Leskinen et al. 2017; Ajuebor et al. 2018; Philipson et al. 2018).

## Materials and Methods

Sample collection, isolation, purification, amplification, and phage characterization followed the HHMI SEA-PHAGES Phage Discovery Guide (https://seaphagesphagediscoveryguide.helpdocsonline.com/home; last accessed 2020/04/30), with modifications indicated below. Library preparation, sequencing, assembly, and gene annotation followed the HHMI SEA-PHAGES Phage Genomics Guide (https://seaphagesbioinformatics.helpdocsonline.com/home; last accessed 2020/04/30).

### Sample Collection and Isolation

To locate phage, ~50 soil samples were collected from various locations in Arizona and plaque assays were performed on the sample filtrates. Most samples did not produce phage that could infect *Staphylococcus* spp. The sample that produced Metroid was collected from a shaded and well-irrigated garden on Arizona State University’s Tempe campus (33.417708N, 111.935974W; ambient temperature 37.7°C). The soil was loosely packed into half of a 15 mL conical tube and stored at 4°C until phage isolation and a plaque assay were performed. In order to isolate bacteriophages, the sample was submerged in 10 mL PYCa liquid media (yeast, tryptone, 1M CaCl_2_, 40% dextrose, cycloheximide), vortexed for one minute, and placed in a shaking incubator at room temperature for 30 minutes. This sample was then centrifuged at 4500 rpm for four minutes and filter-sterilized with a 0.22 μm syringe filter. A 250 μL sample of this filtrate was mixed with 250 μL of host bacteria of the genus *Staphylococcus*, which had been grown to saturation in PYCa and stored at 4°C. After a ten minute incubation at room temperature, the 500 μL of phage plus bacteria was added to 4.5 mL molten PYCa top agar (60°C) and immediately plated on a PYCa agar plate which was incubated for 48 hr at 37°C.

### Purification and Amplification

Smooth plaques appeared on the PYCa plates after 48 hours and were ~3 mm in diameter. One plaque was picked with a sterile pipette tip, and phage were resuspended in phage buffer (1M Tris, 1M MgSO_4_, NaCl, ddH2O, 100 mM CaCl_2_), and a series of six 10-fold serial dilutions were performed. Each dilution was inoculated with 250 μL of *Staphylococcus* spp. host bacteria and incubated at room temperature for ten minutes. Each dilution was plated with 4.5 mL PYCa top agar and incubated at 37°C for 48 hours. A plaque from the plate representing the 10^−2^ dilution was selected to complete two additional rounds of purification through subsequent dilutions and plaque assays. For each purification, we chose to pick plaques from a ‘countable’ plate, on which plaques were separated enough to suggest that each grew from a single phage particle (typically a countable plate had 30 to 300 plaques).

Once purified, we amplified the phage to obtain a titer greater than 1×10^9^ PFU/mL which would provide enough DNA for genome sequencing. A plate containing numerous purified phage plaques was flooded with 8 mL of phage buffer and set at room temperature for an hour to yield a phage lysate. The lysate was collected in a 15 mL tube and centrifuged at 8000 rpm for four minutes then filtered through a 3 mL syringe with a 0.22 μL filter. 10-fold serial dilutions were made with the collected lysate for amplification. A spot titer was made with the undiluted lysate as well as 10^−1^ to 10^−10^ lysate dilutions. Based on counting the number of plaques formed by each lysate in the spot titer assay, the 10^−8^ dilution was selected as the best candidate to produce a countable plate. A full titer plate was prepared with the 10^−7^, 10^−8,^, and 10^−9^ dilutions. The titer calculated from the full titer assay was 2.65×10^10^ PFU/mL.

### Phage Characterization

#### DNA Extraction

DNA extraction was performed on the phage lysate using the Wizard DNA Clean-Up kit (Promega) with minor modifications. 5 μL of nuclease mix (NaCl, ddH_2_O, DNase 1, RNase A, glycerol) was added to 1 μL of lysate and mixed by inversion. The solution was incubated at 37°C for ten minutes. 15 μL of EDTA and 1 μL of Proteinase K were added to the solution and incubated at 37°C for 20 minutes. 2 mL of Wizard DNA Clean-Up resin (Promega) was added to the solution and mixed by inversion for two minutes. The solution was syringed-filtered through two Wizard Genomic DNA columns (Promega) and then washed three times with 80% isopropanol. The columns were twice spun in a centrifuge at top speed for two minutes and then placed in a 90°C heat block for one minute. 50 μL of ddH_2_O was used for elution. Final elutes were combined for 100 μL of total DNA extract. A Nanodrop ND 1000 was used to determine a DNA concentration of 114.9 ng/μL.

#### Transmission Electron Microscopy

A high-titer lysate was made up for Transmission Electron Microscopy (TEM) by spinning 100 μL of phage lysate in a 4°C centrifuge at top speed for 22 minutes. The supernatant was removed and the pellet was resuspended in 10 μL of phage buffer. The high-titer lysate then underwent TEM preparation by negatively staining the virus particles. Specifically, isolated particles were adhered to a 300-mesh carbon-formvar grid for one minute, followed by staining with 1% aqueous uranyl acetate for 30 seconds. Images were acquired using a Philips CM12 TEM operated at 80kV and equipped with a Gatan model 791 CCD camera.

### Library Preparation, Sequencing, and *De Novo* Assembly

A sequencing library was prepared from genomic DNA by using an NEB Ultra II FS kit with dual-indexed barcoding and sequenced on an Illumina MiSeq, yielding a total of 901,246 single-end 150bp reads (>895X coverage). Quality control checks using FastQC v.0.11.7 (http://www.bioinformatics.babraham.ac.uk/projects/fastqc; last accessed 2020/04/30) indicated that the data was of high quality, rendering additional read processing prior to assembly unnecessary. Following Russell (2018), reads were *de novo* assembled using Newbler v2.9, resulting in a single linear contig of size 150,935bp, which was checked for completeness, accuracy, and phage genomic termini using Consed v.29. All software was executed using default settings.

### Genome Annotation

Annotation was performed using DNA Master v.5.23.3 (http://cobamide2.bio.pitt.edu; last accessed 2020/04/30). Putative protein-encoding open reading frames (genes) were identified using Glimmer v.3.0 (Delcher et al. 1999) and GeneMark v.2.5 (Lukashin and Borodovsky 1998) with AUG (methionine), UUG and CUG (leucine), GUG (valine), and AUA (isoleucine) as start codons. Using annotated bacteriophage sequences from public databases, functional assignments were made with Blastp v.2.9 (Altschul et al. 1990), NCBI’s Conserved Domain Database (Marchler-Bauer et al. 2015), and HHPred (Söding et al. 2005). In addition, TMHMM2 (http://www.cbs.dtu.dk/services/TMHMM/; last accessed 2020/04/30) and SOSUI (http://harrier.nagahama-i-bio.ac.jp/sosui/sosui_submit.html; last accessed 2020/04/30) were used to identify membrane proteins. tRNAs were annotated using Aragon v.1.1 (included in DNA Master) and v.1.2.38 (http://130.235.244.92/ARAGORN/; last accessed 2020/04/30) as well as tRNAscan-SE v.2.0 (http://trna.ucsc.edu/tRNAscan-SE/; last accessed 2020/04/30). All software was executed using default settings.

### Comparative genomics analysis

Due to their similar length, number of genes and tRNAs, as well as GC-content, the genomes of the phages IME-SA1, IME-SA2, ISP (Vandersteegen et al. 2011), JA1 (Ajuebor et al. 2018), K (Gill 2014), vB_SauM_0414_108 (Philipson et al. 2018), and vB_SauM-fRuSau02 (Leskinen et al. 2017) were downloaded from GenBank (Table 1) to create a database of *Staphylococcus* cluster C1 phages (Olivera et al. 2019) using PhamDB (https://github.com/jglamine/phamdb/wiki/Using-PhamDB; last accessed 2020/04/30). This custom database was used for all subsequent comparative analyses. First, a multiple sequence alignment was performed utilizing Kalign v.1.04 (Lassmann and Sonnhammer 2005) to produce a neighbor-joining tree. Second, dotplots, comparing the relatedness of different nucleotide sequences, were generated in 10bp sliding windows using Gepard v.1.40 (Krumsiek et al. 2007). Lastly, the database was loaded into Phamerator (https://github.com/scresawn/Phamerator; last accessed 2020/04/30) to visually compare phage genomes.

**Table 1:** Features of the seven *Staphylococcus* cluster C1 phages used for comparative genome analyses.

| name | length | # genes | # tRNAs | GC-content | host | GenBank<br>accession number | reference |
| --- | --- | --- | --- | --- | --- | --- | --- |
| IME-SA1 | 140,218 | 209 | 4 | 30.33 | *S. aureus* | KP687431.1 | unpublished |
| IME-SA2 | 140,906 | 212 | 4 | 30.33 | *S. aureus* | KP687432.1 | unpublished |
| ISP | 138,339 | 215 | 4 | 30.42 | *S. aureus* | FR852584.1 | Vandersteegen et al. 2011 |
| JA1 | 147,135 | 233 | 4 | 30.25 | *S. aureus* | MF405094.1 | Ajuebor et al. 2018 |
| K | 148,317 | 233 | 4 | 30.39 | *S. aureus* | NC_005880.2 | Gill 2014 |
| vB_SauM_0414_108 | 151,627 | 249 | 4 | 30.39 | *S. aureus* | MH107769.1 | Philipson et al. 2018 |
| vB_SauM-fRuSau02 | 148,464 | 236 | 4 | 30.22 | *S. aureus* | MF398190.1 | Leskinen et al. 2017 |

## Results and Discussion

The complete genome sequence of the *Staphylococcus* bacteriophage Metroid was sequenced and annotated (see “Materials and Methods” for details). The *Myoviridae* morphology (*i.e.*, an icosahedral capsid [diameter: 100nm] enclosing the double-stranded DNA attached to a long, contractile tail [length: 108nm]; Figure 1a) as well as the genome size of 151kb (including the ~10kb terminal repeat) suggests that Metroid belongs to the *Staphylococcus* phage cluster C. Metroid’s genome has a GC-content of 30.40%, similar to those of previously published *Staphylococcus* phages (27.98-34.96%) (Kwan et al. 2005; Deghorain et al. 2012; Olivera et al. 2019). The tightly-packed genome contains 254 predicted protein-coding genes as well as 4 tRNAs, most of which are transcribed on the forward strand (Figure 1b). This corresponds to a gene density of 1.68 genes/kb – on the upper end of the range previously reported for cluster C phages (164-249 genes; 0-5 tRNAs; 1.25-1.64 genes/kb) (Olivera et al. 2019). Although the overall gene coding potential of Metroid is 89.42%, only 26 of the 254 predicted proteins could be assigned a putative function. The majority of predicted proteins are either conserved but of no known function (170 out of 254), membrane proteins (22), or unique (*i.e.*, without a match to any of the queried databases; 1). Thereby, functionally related genes are organized into distinct modules (*e.g.*, distinct head and tail modules connected by a head-to-tail adapter) (Figure 1b).

**Figure 1:**
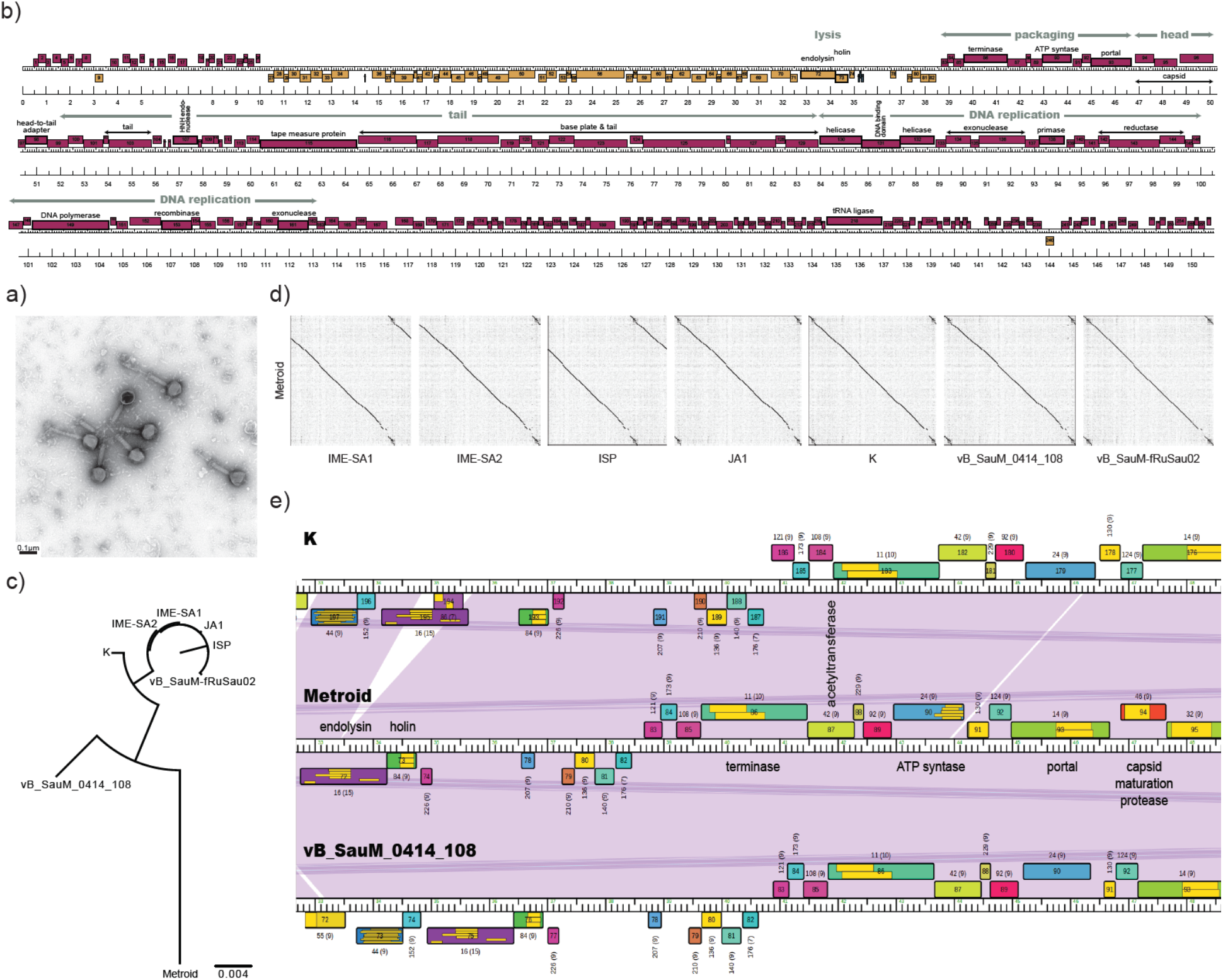
Characterization of Metroid and its relatedness to other *Staphylococcus* cluster C1 phages. a) Transmission electron microscopy image showing Metroid’s morphology. b) Metroid’s genome contains 254 predicted protein-coding genes as well as 4 tRNAs; total genome size: 151kb including the ~10kb terminal repeat. The majority of genes are transcribed on the forward strand as shown in pink; genes transcribed on the reverse strand are highlighted in orange; tRNAs in blue. Functionally related genes are organized into distinct modules (highlighted in grey). c) Neighbor-joining tree and d) dotplot of Metroid and seven previously described *Staphylococcus* bacteriophages (Table 1). e) Genes in the lysis cassettes as well as in the packaging module show a strong conservation between Metroid and two closely-related *Staphylococcus* phages, K (Gill 2014) and vB_SauM_0414_108 (Philipson et al. 2018). Genes are labelled with their putative function, with genes belonging to the same protein family (pham) depicted in the same color. Purple coloring between genomes highlights regions of high nucleotide similarity (i.e., a BLAST e-value of 0).

Comparative genomic analysis with seven *Staphylococcus* subcluster C1 phages indicates that Metroid is most closely related to vB_SauM_0414_108 (Figure 1c,d) – a phage discovered as part of a recent effort proposing a guideline and standardized workflow to submit phages to the Federal Drug Administration to be considered as potential future treatments of bacterial infections (Philipson et al. 2018). More generally, genes in the lysis cassettes show a strong conservation between Metroid and the closely-related *Staphylococcus* cluster C phages, previously characterized for therapeutic research (Figure 1e), suggesting that Metroid might be a suitable candidate for future phage therapies.

## Data availability

Metroid’s genome assembly is available in GenBank under accession number XXXXXX.

## Acknowledgements

This study was supported by the Howard Hughes Medical Institute SEA-PHAGES program and Arizona State University’s School of Life Sciences. DNA concentration was determined in the Arizona State University DNA Shared Resource Facility. Library preparation and sequencing was performed at the University of Pittsburgh. Computations were partially performed at Arizona State University’s High Performance Computing facility. We are grateful to David Lowry for transmission electron microscopy imaging, Billy Biederman, Graham Hatfull, Deborah Jacobs-Sera, Welkin Pope, Daniel Russell, and Vic Sivanathan for library preparation, sequencing, and assembly as well as providing faculty training for the SEA-PHAGES program.

